# Local recruitment of DNA repair proteins enhances CRISPR-ssODN-HDR editing

**DOI:** 10.1101/2022.04.13.488255

**Authors:** Nathaniel Jillette, Jacqueline J. Zhu, Albert W. Cheng

## Abstract

CRISPR-Cas technologies enable precise editing of genomic sequences. One major way to introduce precise editing is through homology directed repair (HDR) of DNA double strand breaks (DSB) templated by exogenously supplied single-stranded oligodeoxyribonucleotides (ssODN). Competing pathways determine the outcome of edits. Non-homologous end-joining pathways produce destructive insertions/deletions (indels) at target sites and are dominant over the precise homology directed repair pathways. In this study, we aim to favor HDR and use two strategies to recruit DNA repair proteins (DRPs) to Cas9 cut site, the Casilio-DRP approach that recruits RNA-binding protein-tethered DRPs to target site via aptamers appended to guide RNA; and the 53BP1-DRP approach that recruits DRPs to DSBs via DSB-sensing activity of 53BP1. We conducted two screens using these approaches and identified DRPs such as FANCF and BRCA1 that when recruited to Cas9 cut site, enhance ssODN-templated HDR and increase the proportion of precise edits. This study provides not only new constructs for enhanced CRISPR-ssODN-HDR but also a collection of DRP fusions for studying DNA repair processes.

## Introduction

The development of programmable nucleases has opened an avenue for editing genome sequences of virtually any cell or organism, transforming basic and medical research, and enabling applications from agriculture to therapeutics [1, 2]. However, realizing the full potential of this revolutionary technology in patients will depend on highly efficient and specific editing of the genome with nucleotide precision and in sufficient numbers of cells to have therapeutic value. Programmable nucleases introduce a double stranded break (DSB) at a desired site in the genome whereupon the DSB may be repaired by one of a variety of cellular DNA repair mechanisms. The most efficient repair pathway is error-prone non-homologous end joining (NHEJ), which has been widely exploited to disrupt gene function through the introduction of random insertions/deletions (indels) [3]. However, to introduce precise changes that could be used to correct disease-causing variants, exogenous DNA templates with homology to the target region are required to insert new DNA sequences at the site of the DSB. The cellular machinery used for homology-directed repair (HDR) depends on the repair template itself: whereas double-stranded supercoiled donor plasmids can undergo classical homologous recombination (HR) with the genome, the mechanism of repair mediated by single-stranded DNA templates is not completely understood and several pathways have been proposed [4-11]. NHEJ and HDR represent competing pathways that contain overlapping yet distinct protein components. NHEJ is the dominant outcome even in the presence of a repair template. Attempts to block NHEJ genetically or with protein and chemical inhibitors shift the balance in favor of HDR [12-15] but may risk unwanted genome instability. The alternative approach is to stimulate the HDR pathway. Indeed, modest improvement in HDR efficiency has come from the overexpression/activation of proteins specific to the HDR pathway such as RAD51 [16] or the fusion of DNA repair proteins with Cas9 nuclease [17]. Other orthogonal strategies that have been reported to slightly improve HDR frequencies include the delivery of recombinant Cas9 ribonucleoprotein [18, 19], cell cycle control of Cas9 expression or perturbing the cell cycle [20, 21], cold shock [22] and tethering the repair template to the nuclease [23]. Yet despite these advances, NHEJ remains the predominant repair event. Thus, with the added challenge of delivering both the nuclease and the repair template to cells *in vivo*, it is imperative to dramatically improve HDR efficiency such that it is the preferred repair outcome in treated cells.

We previously developed a hybrid system based on CRISPR/Cas9 and the programmable Pumilio RNA-binding protein, termed “Casilio,” to recruit effector proteins at genomic targets [25]. Here, we employ Casilio to recruit DNA repair proteins or their complexes at the nuclease target site to favor HDR of DSB. As an alternative strategy, we fused DNA repair proteins to the DSB-sensing domain of p53 binding protein 1 [26] so that they are actively recruited to Cas9-generated DSB. We identify several DRPs, and their combinations, when recruited to the Cas9 target site or Cas9-generated DSBs, increase repair towards the HDR pathway. In addition to providing a new strategy to enhance precise editing, the collection of Casilio-tethered or DSB-targeting DRP modules will be useful for studying DNA repair processes.

## Results

### Screening a library of Casilio-tethered DRPs for HDR enhancement

For high-throughput quantification of genome editing outcomes by flow cytometry, we have constructed an HDR/NHEJ reporter HEK293T cell line with a constitutively expressed BFP transgene inserted into the AAVS1 locus. Editing experiments can be performed by co-delivering Cas9 and a sgRNA targeting BFP with a single-stranded oligodeoxyribonucleotides (ssODN) that changes BFP to GFP by incorporating H66Y mutation. This reporter system conveniently reports on the fraction of cells that have undergone HDR (GFP+ population) or NHEJ (GFP-/BFP-double negative population). We identified 78 DRPs and mutants of interest including many previously implicated in HDR or single-strand DNA templated repair processes. We fused DRPs individually to N- or C-terminal of PUFa to generate a Casilio-DRP library with 140 constructs (**Supplementary table 1**) that can be individually tested for HDR enhancement, using the HDR/NHEJ reporter cell line (**Fig. 1a**). We transfected Cas9, sgRNA targeting BFP with 5 copies of PBSa (sgBFP-5xPBSa), and each of the Casilio-DRP constructs into HEK293T HDR/NHEJ cell line, and conduct flow cytometry on Day 7. HDR/NHEJ ratio was calculated by dividing the percentage of GFP+ cells by the percentage of BFP-/GFP-cells. To allow comparison between different batches of experiments, we normalized the HDR/NHEJ to the empty vector control by dividing the test HDR/NHEJ ratio by the control HDR/NHEJ ratio to obtain the fold improvement of HDR/NHEJ ratio (hereafter referred to as fold improvement, FI). From the screen, we identified 23 constructs with FI≥1.5, namely, FANCF (both termini), BRCA1 (both termini), RAD54L-PUFa, PALB2-(WT, KR-mutant, delETGE mutant)-PUFa, LIG1 (both termini), RECQ5-(Full length, strand annealing (SA) domain)-PUFa, FEN1 (both termini), FANCG (both termini), PUFa-FANCM, CtIP-PUFa, PUFa-FANCP, XRCC3 (both termini), and RAD51-PUFa, with FANCF and BRCA1 ranking on top (**Fig 1b,c**).

**Fig. 1.**
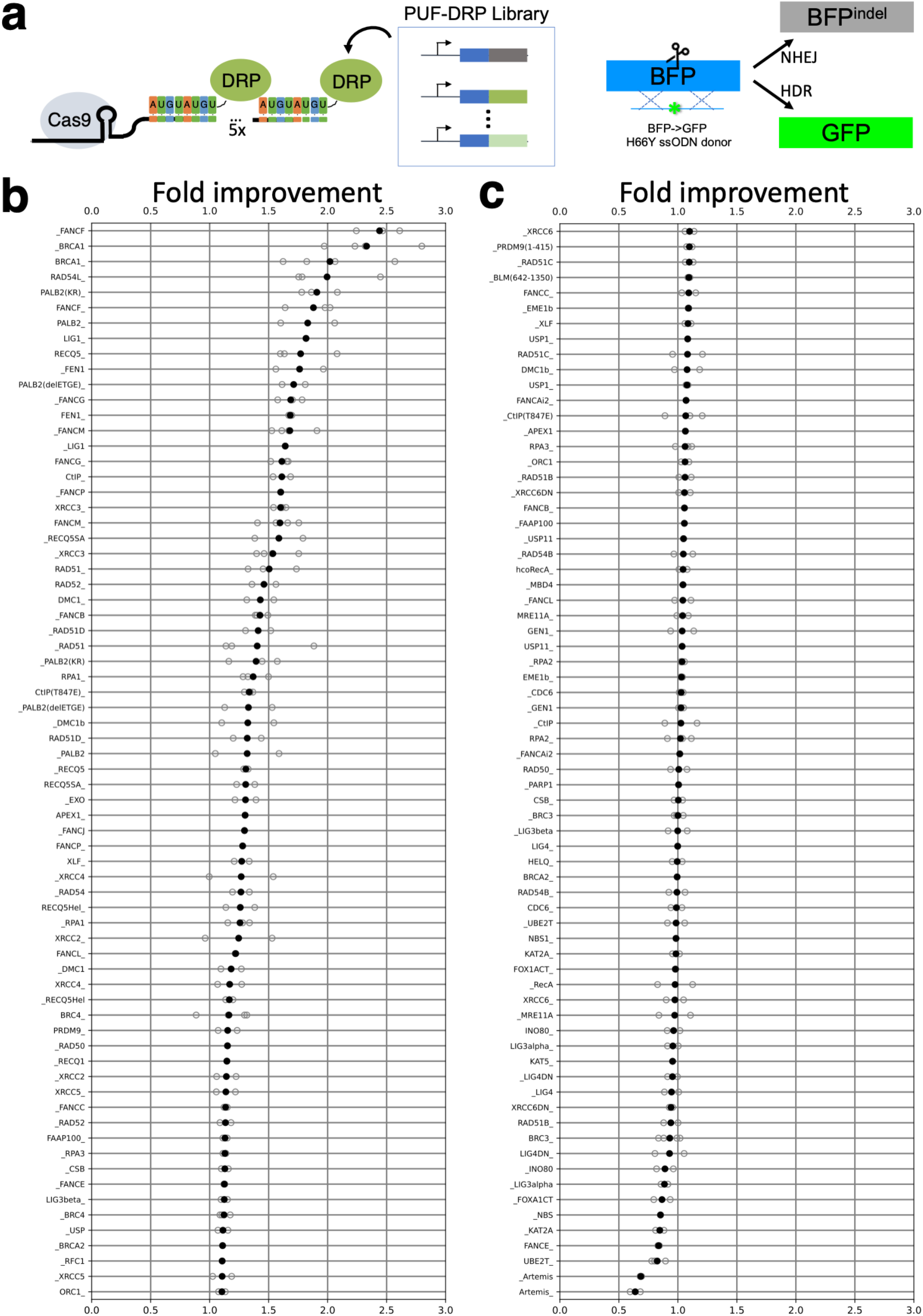
Screening of Casilio-DRP library for HDR enhancement. **(a**) A Casilio-DRP library consists of 140 constructs derived from 78 DNA repair proteins (DRPs) or mutants individually fused to N- or C-termini of PUFa that can be recruited by target-specific sgRNA (in this case, sgBFP targeting BFP ORF) appended with 5 copies of binding sites for PUFa (5xPBSa). Plasmids encoding Cas9, sgBFP-5xPBSa, BFP->GFP ssODN donor encoding the H66Y mutation and individual PUFa-DRP fusions were co-transfected into a HEK293T HDR/NHEJ reporter cell line carrying a CAGGS promoter-driven BFP transgene at the AAVS1 locus. NHEJ repair after Cas9 cutting generated BFP with indels leading to loss of fluorescence while HDR using ssODN as template results in GFP expression. HDR and NHEJ events were derived from %GFP+ and %BFP-GFP-cells measured by flow cytometry, respectively and are used to derive HDR/NHEJ ratio. To allow comparison between batches of experiments, the HDR/NHEJ ratio of each factor is normalized to that of the sample transfected with empty vector (EV) to obtain fold improvement (FI). **(b**,**c)** A plot of FI values of the first 70 (b), and the last 70 (c) Casilio-DRP constructs ranked by mean FI. DRP_ and _DRP denote N- and C-terminal fusions, i.e., DRP_PUFa, PUFa_DRP, respectively. Hollow circles denote values from replicate experiments. Filled circles denote mean values, or the value for constructs screened only once.

### Screening a library of 53BP1-DRPs for HDR enhancement

As an alternative strategy, we fused DNA repair proteins to double strand break (DSB)-sensing domain of p53 binding protein 1 (amino acids 1220-1714 of 53BP1, hereafter referred simply as 53BP1). We generated a 53BP1-DRP library with 59 constructs derived from 51 DRPs and mutants fused to N- or C-termini of 53BP1 (**Supplementary Table 2**). We co-transfected Cas9, a sgRNA targeting BFP (sgBFP), and individual 53BP1-DRP modules to HEK293T HDR/NHEJ cell line, and calculated FI of HDR/NHEJ (**Fig. 2a**). Again, BRCA1 and FANCF were recovered as the top HDR enhancers (**Fig. 2b**). In addition, XRCC3 and CtIP(T847E) were third and forth in the list, among a total of 17 with FI≥1.5.

**Fig. 2.**
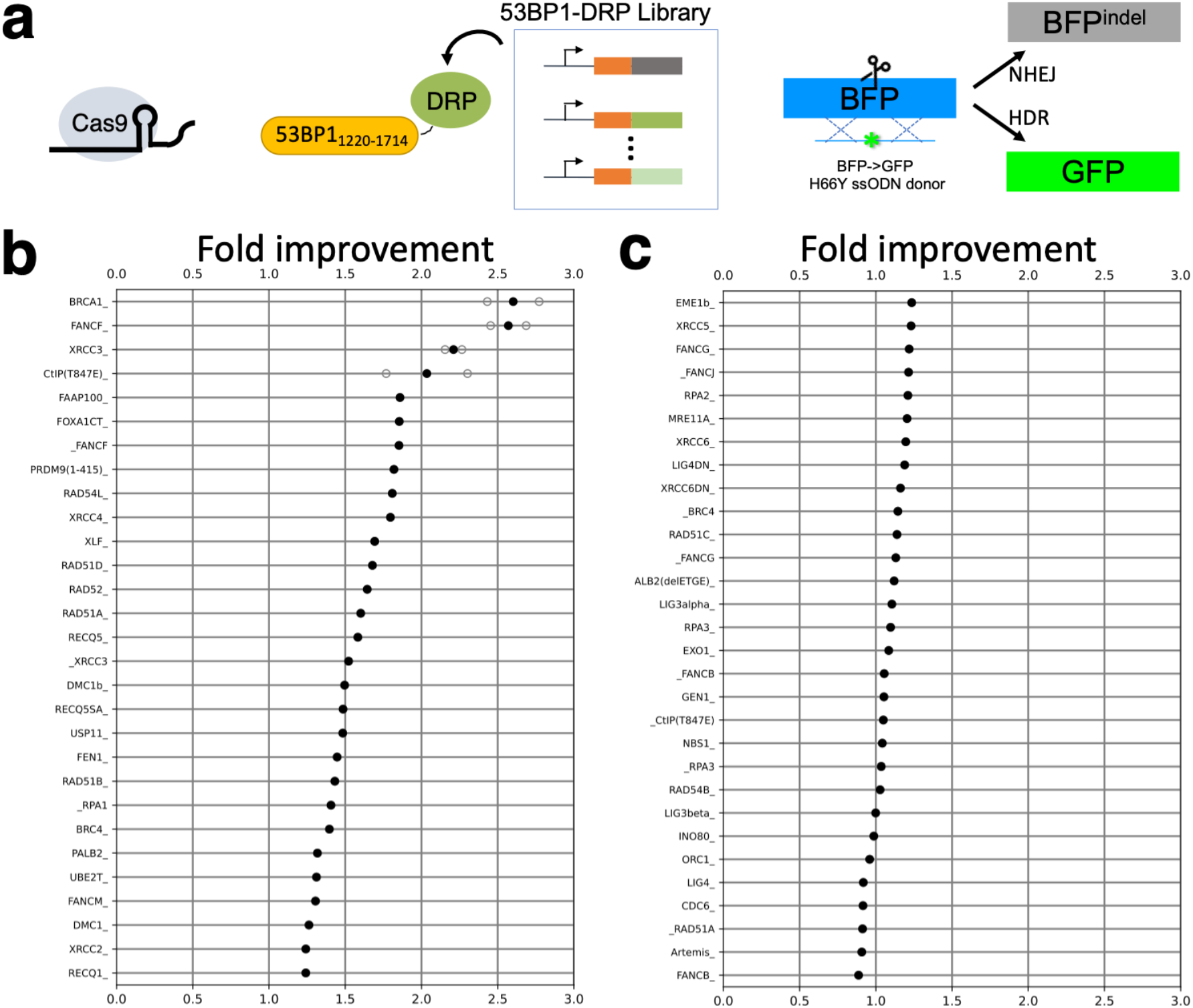
Screening of 53BP1-DRP library for HDR enhancement. **(a**) A 53BP1-DRP library consists of 51 DNA repair proteins (DRPs) or mutants individually fused to N- or C-termini of 53BP1 that are recruited to double strand break sites (DSBs) generated by Cas9. Plasmids encoding Cas9, sgBFP, BFP->GFP ssODN donor encoding the H66Y mutation and individual 53BP1-DRP were co-transfected into a HEK293T HDR/NHEJ reporter cell line carrying a CAGGS promoter-driven BFP transgene at the AAVS1 locus. NHEJ repair after Cas9 cutting generated BFP with indels leading to loss of fluorescence while HDR using ssODN as template results in GFP expression. HDR and NHEJ events were derived from %GFP+ and %BFP-GFP-cells measured by flow cytometry, respectively and are used to derive HDR/NHEJ ratio. To allow comparison between batches of experiments, the HDR/NHEJ ratio of each factor is normalized to that of the sample transfected with empty vector (EV) to obtain fold improvement (FI). **(b**,**c)** A plot of FI values of the first 29 (b), and the last 30 (c) 53BP1-DRP constructs ranked by mean FI. DRP_ and _DRP denote N- and C-terminal fusions, i.e., DRP_53BP1, 53BP1_DRP, respectively. Hollow circles denote values from replicate experiments. Filled circles denote mean values or the value for the constructs screened only once.

### Enhancement of HDR by Casilio-DRPs and 53BP1-DRPs at endogenous genes

We next investigated whether HDR-enhancing constructs identified from the reporter screens could be applied to enhancing the editing of endogenous genes. For 53BP1-DRP system, we selected FANCF, BRCA1, CtIP(T847E) and XRCC3 and tested either FANCF-53BP1 alone or a mix of the four constructs (**Fig 3a**). We selected two popular test genes *EMX1* and *DYRK1* and used previously designed sgRNA and ssODN [21] (**Fig 3b,c**). We co-transfected Cas9, sgEMX1 or sgDYRK1, and FANCF-53BP1 or the mix of four constructs (4F-53BP1), and extracted genomic DNA on Day 3 post-transfection. Target-specific primers were used to amplify regions surround the editing site which were then sent for Sanger sequencing. Sequencing traces were analyzed by TIDER [27] to obtain percentages of sequences with signature of either HDR or NHEJ. While about 40% to 50% of edited sequences present NHEJ outcomes in non-DRP-supplemented Cas9 samples for *EMX1* (HDR/NHEJ=1.08) and *DYRK1* (HDR/NHEJ=0.68), the addition of FANCF-53BP1 construct switched major editing outcome to HDR, increasing HDR/NHEJ to 2.93 (FI=2.71) and 1.31 (FI=1.93) for *EMX1* and *DYRK1*, respectively. The addition of other three constructs to a four construct cocktail (4F-53BP1) further increased HDR/NHEJ to 6.2 (FI=5.73) and 2.3 (FI=3.38) for *EMX1* and *DYRK1*, respectively (**Fig 3d,e**). We next tested Casilio-DRP, with FANCF-PUFa alone or with additional BRCA1, CtIP(T847E) and XRCC3 (4F-PUFa) (**Fig 3f**). FANCF-PUFa increased HDR/NHEJ from 1.32 to 2.96 (FI=2.24) for *EMX1*, and from 0.25 to 0.88 for *DYRK1* (FI=3.51). The four-construct combination tethered by PUFa further increased HDR/NHEJ to 3.54 (FI=2.69) and 1.67 (FI=6.63), for *EMX1* and *DYRK1*, respectively. These results together confirmed that the Casilio-DRP and 53BP1-DRP provides HDR enhancement activity at endogenous genes.

**Fig. 3.**
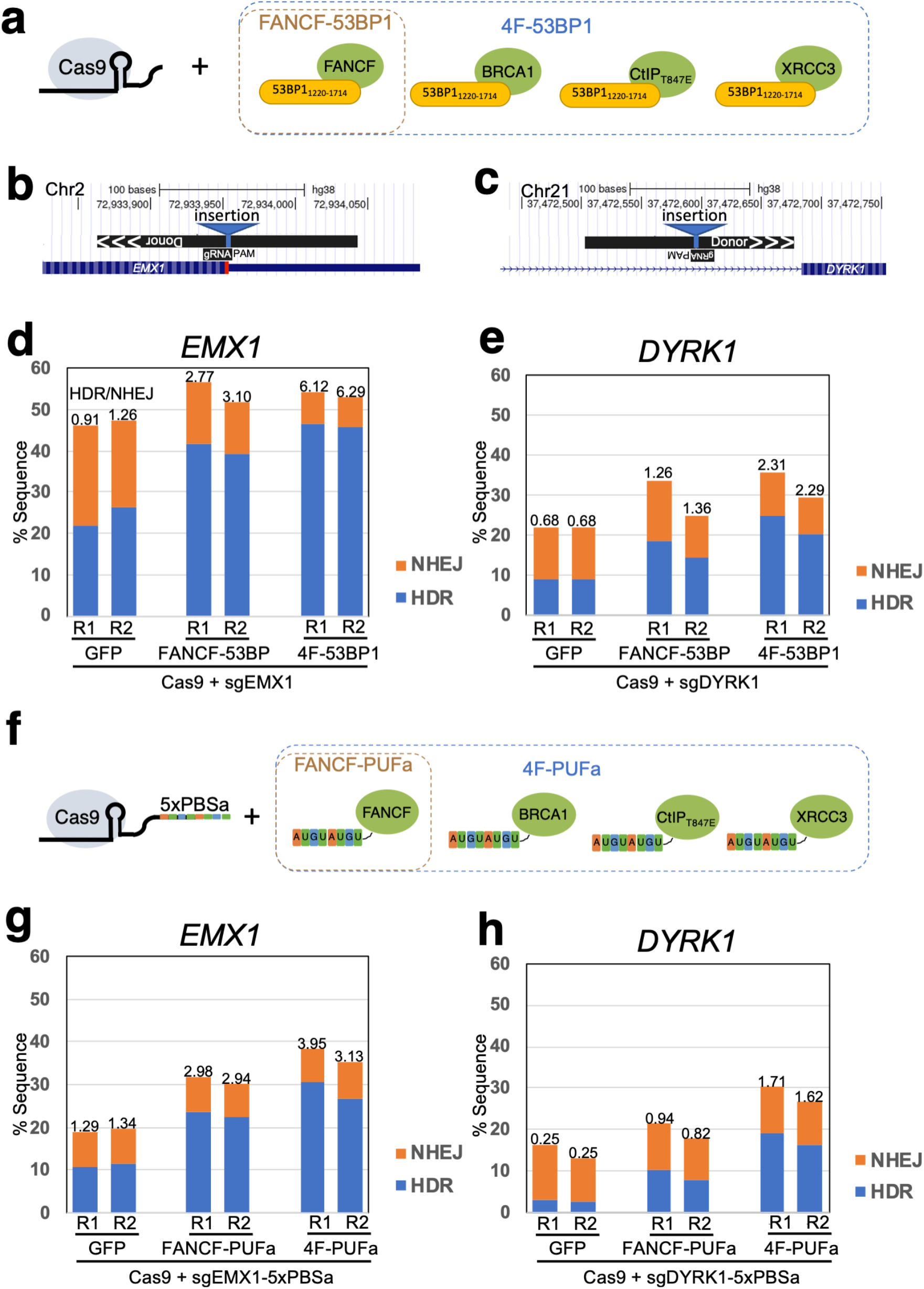
HDR enhancement by Casilio-DRPs and 53BP1-DRPs at endogenous target genes. **(a**) Schematic of tested DRP-53BP1 combinations (FANCF-53BP1 alone or 4F-53BP1, a mix of FANCF, BRCA1, CtIP(T847E), XRCC3 tethered by 53BP1). **(b**,**c)** Annotation of ssODNs and sgRNAs with respect to the target genes *EMX1* (b) and *DYRK1* (c). **(d**,**e)** Column plots of percentage of *EMX1* (d) and *DYRK1* (e) amplicons with sequence signature resulting from NHEJ (red) and HDR (blue) for replicate 1 (R1) and replicate 2 (R2) experiments transfected with the indicated constructs. 4F-53BP1 indicate a mixture of FANCF, BRCA1, CtIP(T847E), XRCC3 tethered by 53BP1. HDR/NHEJ ratio indicated on top of columns **(f)** Schematic of tested DRP-PUFa combinations (FANCF-PUFa alone or 4F-PUFa, a mix of FANCF, BRCA1, CtIP(T847E), XRCC3 tethered by PUFa). **(g**,**h)** Column plots of percentage of *EMX1* (g) and *DYRK1* (h) amplicons with sequence signature resulting from NHEJ (red) and HDR (blue) for replicate 1 (R1) and replicate 2 (R2) experiments transfected with the indicated constructs. 4F-PUFa indicate a mixture of FANCF, BRCA1, CtIP(T847E), XRCC3 tethered by PUFa. HDR/NHEJ ratio indicated on top of columns.

## Discussion

In this study, we evaluated two strategies for recruiting DNA repair proteins (DRPs) to Cas9 cut site to enhance HDR templated by single stranded oligodeoxyribonucleotide (ssODN). The Casilio approach uses extended gRNA with pumilio binding sites (PBS) to recruit DRP fused to cognate pumilio/PUF domain. The 53BP1 approach tethers DRPs to 53BP1 double strand break (DSB)-sensing domain to recruit DRPs to DSBs generated by Cas9. We generated a Casilio-DRP library with 140 constructs for 78 DRPs, and a 53BP1-DRP library with 59 constructs for 51 DRPs. To facilitate screening of DRP constructs for HDR enhancement activity, we generated a HEK293T cell line with AAVS-integrated CAGGS-driven BFP transgene to act as HDR/NHEJ reporter when used with an ssODN encoding H66Y mutation changing BFP to GFP upon successful HDR and destroying BFP signal in case of NHEJ. In the Casilio-DRP and 53BP1-DRP screens, we recovered 23 (out of 140) and 17 (out of 59) constructs with HDR-enhancing activity with ≥1.5 fold improvement (FI) compared to control. Several factors were identified on top in both screens, namely FANCF, BRCA1, CtIP, and XRCC3. We next tested these constructs on two endogenous genes, *EMX1* and *DYRK1*. The FANCF-tethering constructs produced improved HDR with up to 3.5 FI while four factor combination with FANCF, BRCA1, CtIP(T847E) and XRCC3 provided up to ∼6.6 FI. In this present study, we only worked on Cas9-induced DSBs repaired by ssODN, while other template types (circular double strand plasmids, double strand linear DNA, etc.), DNA lesion types (DSB, single strand nicks) may benefit from different DRPs. In future studies, screens with these different parameters should be conducted with our collection of DRP fusion constructs. We believe that we not only provide new modules for enhanced CRISPR editing but also two libraries of useful DRP fusions applicable to the study of DNA repair processes.

## Methods

### Cloning

Coding sequences of DNA repair proteins (DRPs) were amplified from cDNA library extracted from HEK293T, then cloned into pCR8GWTOPO-based backbone encoding PUFa or 53BP1(1220-1714) with SgrAI-AclI or FseI-PacI cloning site pairs for N- and C-terminal insertion of DRPs, respectively. The resultant pCR8GWTOPO plasmids containing the Casilio-DRP or 53BP1-DRP ORFs were then transferred to pAC90-pmaxDEST vector (Addgene Plasmid #48222) or pcDNA3.1-DEST by LR Clonase II reaction (Invitrogen, ThermoFisher Scientific #11791100) for Mammalian expression.

Guide RNA are under control of the human U6 promoter. gRNA spacer sequences were cloned into sgRNA-PBS or sgRNA expression vectors via an oligonucleotide-annealing protocol [28].

### Guide RNA target sequences and ssODN sequences

sgBFP: GCTGAAGCACTGCACGCCAT

sgEMX1: GTCACCTCCAATGACTAGGG

sgDYRK1: GTTCCTTAAATAAGAACTTT

BFP->GFP H66Y donor: GCCACCTACGGCAAGCTGACCCTGAAGTTCATCTGCACCACCGGCAAGCTGCCCGTGCCCTGGCCCACCCTCGTGACCACCCTGACGTACGGCGTGCAGTGCTTCAGCCGCTACCCCGACCACATGA

EMX1-donor (insertion lowercase underlined): GGGAGTGGCCAGAGTCCAGCTTGGGCCCACGCAGGGGCCTGGCCAGCAGCAAGCAG CACTCTGCCCTCGTGGGTTTGTGGTTGCCCACCC<u>ggatccaagctt</u>TAGTCATTGGAGGTGA CATCGATGTCCTCCCCATTGGCCTGCTTCGTGGCAATGCGCCACCGGTTGATGTGATGGGAGCCCTTCTTCTTC

DYRK1-donor (insertion lowercase underlined): ATCTTCTATACCATTAAAAACATAACTGTTGTGTTGAGTAACATATACCTGTTTGTAG TTAGAAAAGTTTTTTAATATTGAATATCCTAAA<u>gctagcaagct</u>TTCTTATTTAAGGAACCA TTAGATATGTCAAATGATACAAACATTAGGATATGAATATTTCCTTTAAACCTCACTTATCTTC

### Cell culture

HEK293T (ATCC # CRL-3216) were cultivated in Dulbecco’s Modified Eagle’s Medium (DMEM) (Sigma #D5671) with 10% fetal bovine serum (Gibco, ThermoFisher Scientific #10437028), 4% Glutamax (Gibco, ThermoFisher Scientific #35050-061), 1% Sodium Pyruvate (Gibco, ThermoFisher Scientific #11360-070) and 1% penicillin-streptomycin (Gibco, ThermoFisher Scientific #15140-163).

### Transfection

HEK293T HDR/NHEJ cell line was generated by transfecting cells with Cas9, AAVS1 targeting sgRNA, and a knock-in construct containing homology arms surrounding the cut site as well as 2A-puro and CAGGS-BFP. For Casilio-DRP screen, HEK293T HDR/NHEJ cells were seeded at a density of 100,000 cells/well in a 12-well cell culture plate 24 hours before transfection. Cells were then transfected with 200ng of ssODN encoding BFP->GFP mutation, 133 ng of Cas9 plasmid, 133ng of sgBFP-5xPBSa plasmid, and 133ng of PUFa-DRP plasmid using 1.5 µl Lipofectamine 3000 (Invitrogen, ThermoFisher Scientific #L3000075). For 53BP1-DRP screen, cells were seeded and transfected similarly but with sgBFP-5xPBSa substituted by sgBFP plasmid, and with 53BP1-DRP constructs instead of PUFa-DRP constructs. For endogenous gene editing experiments with 53BP1-DRP, cells were transfected with 200ng ssODN, 200ng pX330-GFP vector carrying both Cas9 and sgRNA, and 200ng FANCF-53BP1 plasmid or 50ng each of FANCF-53BP1, BRCA1-53BP1, CtIP(T847E)-53BP1 and XRCC3-53BP1 plasmids. For endogenous gene editing experiments with Casilio-DRP, cells were transfected with 200ng ssODN, 100ng Cas9 plasmid, 100ng sgRNA-5xPBSa plasmid, and 200ng FANCF-PUFa plasmid or 50ng each of FANCF-PUFa, BRCA1-PUFa, CtIP(T847E)-PUFa, and XRCC3-PUFa plasmids. Media was changed at 24 hours post-transfection.

### Flow cytometry

For estimation of HDR and NHEJ events, transfected cells were passaged once on Day 3 by splitting cells at ∼1:10. On Day 7, cells were trypsinized, suspended in media then analyzed on a LSRFortessa X-20 or FACSymphony flow cytometer (BD Bioscience). Fifty thousand events were collected each run.

### PCR-based assay for HDR/NHEJ determination

For quantification of HDR/NHEJ at endogenous genes, transfected cells were harvested on Day 3 and genomic DNA was extracted using DNeasy Blood & Tissue Kit (QIAGEN # 69506). The region surrounding *EMX1* target site was amplified with EMX1-F: GCCATCCCCTTCTGTGAATGTTAGAC and EMX1-R: GGAGATTGGAGACACGGAGAGCAG primers. The region surrounding *DYRK1* target site was amplified with DYRK1-F: GAGGAGCTGGTCTGTTGGAGAAGTC and DYRK1-R: CCCAATCCATAATCCCACGTTGCATG primers. The amplicons were then submitted for Sanger sequencing. Sequencing traces were analyzed using TIDER at http://shinyapps.datacurators.nl/tider/

## Acknowledgments

This work is partially supported by internal funds provided by the Jackson Laboratory, Arizona State University as well as grants from the National Human Genome Research Institute (R01-HG009900 to A.W.C.). We would like to thank the Flow Cytometry at the Jackson Laboratory for providing the equipment and support for our flow cytometry experiments.

